# Resting-state functional connectivity, cortical GABA and neuroactive steroids in peripartum and peripartum depressed women: a functional magnetic imaging and resonance study

**DOI:** 10.1101/411405

**Authors:** Kristina M. Deligiannidis, Christina L. Fales, Aimee R. Kroll-Desrosiers, Scott A. Shaffer, Vanessa Villamarin, Yanglan Tan, Janet E. Hall, Blaise B. Frederick, Elif M. Sikoglu, Richard A. Edden, Anthony J. Rothschild, Constance M. Moore

## Abstract

Postpartum depression (PPD) is associated with abnormalities in resting-state functional connectivity (RSFC) but the underlying neurochemistry is unclear. We hypothesized that peripartum GABAergic neuroactive steroids (NAS) are related to cortical GABA concentrations and RSFC in PPD as compared to healthy comparison women (HCW). To test this, we measured RSFC with fMRI and GABA+/Creatine (Cr) concentrations with proton magnetic resonance spectroscopy (^1^H MRS) in the pregenual anterior cingulate (pgACC) and occipital cortices (OCC) and quantified peripartum plasma NAS. We examined between-group differences in RSFC and the relationship between cortical GABA+/Cr concentrations with RSFC. We investigated the relationship between NAS, RSFC and cortical GABA+/Cr concentrations. Within the default mode network (DMN) an area of the dorsomedial prefrontal cortex (DMPFC) had greater connectivity with the rest of the DMN in PPD (peak voxel: MNI coordinates (2, 58, 32), p=0.002) and was correlated to depression scores (peak HAM-D17 voxel: MNI coordinates (0, 60, 34), p=0.008). pgACC GABA+/Cr correlated positively with DMPFC RSFC in a region spanning the right anterior/posterior insula and right temporal pole (r=+0.661, p=0.000). OCC GABA+/Cr correlated positively with regions spanning both amygdalae (right amygdala: r=+0.522, p=0.000; left amygdala: r=+0.651, p=0.000) as well as superior parietal areas. Plasma allopregnanolone was higher in PPD (p=0.03) and positively correlated with intra DMPFC connectivity (r=+0.548, p=0.000) but not GABA+/Cr. These results provide initial evidence that PPD is associated with altered DMN connectivity; cortical GABA+/Cr concentrations are associated with postpartum RSFC and allopregnanolone is associated with postpartum intra-DMPFC connectivity.

## INTRODUCTION

Peripartum depressive illnesses affect approximately 1 in 8 women [1], with undetected or undertreated illness associated with negative effects on the woman [2] and her family[3]. Peripartum depression (PPD) is a heterogeneous disorder including a range in symptom severity [4]. Antepartum depressive or anxiety symptoms are a risk factor for PPD [5] and may represent an early manifestation of the disorder. Additional risk factors associated with PPD include a prior depressive episode and prior PPD [6]. PPD pathophysiology is insufficiently understood.

Studies have established network dysfunction in non-peripartum major depressive disorder (MDD)[7]. fMRI studies using a small number of predefined regions point to reduced functional connectivity between regions of the default mode (DMN)[8] and salience networks (SN)[9] in PPD[10]. The DMN is a network of connected brain regions most active at rest and involved in monitoring of the external environment and internal mentation. Although studies differ, this network often includes the medial prefrontal cortex (MPFC), posterior cingulate cortex, precuneus and inferior parietal lobule[11]. The SN is a paralimbic-limbic network active both at rest and during task-related activity which integrates sensory, emotional and cognitive information to contribute to social behavior and self-awareness. The SN often includes the dorsal ACC, anterior insula, amygdala, ventral striatum, dorsomedial thalamus, hypothalamus and the substantia nigra/ventral tegmental area [12]

Γ-aminobutyric acid (GABA) is believed to contribute to intrinsic functional connectivity of a network[13-15] and has been implicated in the pathogenesis of major depression[16] and premenstrual dysphoric disorder[17]. A single study measured occipital cortex (OCC) GABA magnetic resonance spectroscopy (MRS) concentrations in PPD[18] and reported no difference between women with PPD and healthy postpartum women, in contrast to findings in MDD[16]. GABA may modulate activity synchrony between brain regions and may be specific to connectivity within the network in which GABA is measured. For example, a study in healthy subjects reported a negative correlation between GABA concentrations measured by magnetic resonance spectroscopy (MRS) and functional connectivity in the DMN[13]. Task-based fMRI studies in healthy subjects reported a correlation between cortical GABA concentration and OCC activity[19], between cortical GABA and insula activity[14] and between ACC GABA and the ACC activity[15], though not all studies agree[20].

Neuroactive steroids (NAS) are allosteric modulators of GABAergic function[21] and may be associated with altered functional connectivity across the menstrual cycle[22] and in premenstrual dysphoric disorder[23]. Plasma and central nervous system NAS concentrations increase across pregnancy and fall precipitously at delivery [24,25]. Dysregulation of NAS metabolism, including allopregnanolone and pregnanolone, and/or their interaction with GABA, has been implicated in PPD[18,25,26].

We hypothesized that peripartum GABAergic NAS concentrations are correlated to cortical GABA concentrations and RSFC in PPD as compared to healthy women. The study rationale is that NAS modulate GABAergic function[21]; GABA can modulate activity synchrony between brain regions[13] and that altered GABA can lead to circuit dysfunction[27]. This study used an integrated approach to measure group RSFC, cortical GABA+/Creatine (GABA+/Cr) concentrations and peripartum plasma NAS in women who developed PPD during pregnancy up to 8 weeks postpartum as compared to healthy peripartum women. Our main aims were to1) examine RSFC networks with nodes in the ACC/MPFC due to previous reports of altered connectivity of regions within the DMN and SN[10]; 2) measure cortical GABA+/Cr in the pregenual (pg)ACC/MPFC given reduced pgACC GABA in MDD[28] 3) examine associations between pgACC GABA+/Cr and RSFC; and 4) measure peripartum NAS to examine their relationship with PPD, patterns of RSFC and pgACC GABA+/Cr. A validation aim was to measure GABA+/Cr in the OCC, given its use as a region of interest in MDD MRS studies[16] and to compare to the one previous PPD GABA MRS study[18].

We tested the hypotheses: (1) PPD would be associated with altered connectivity in networks with nodes in the ACC/MPFC, 2) PPD would be associated with lower pgACC GABA+/Cr, 3) RSFC of networks with nodes in the ACC/MPFC would be associated with pgACC GABA+/Cr in postpartum women and 4) plasma allopregnanolone, but not pregnanolone, concentrations would be lower in PPD and associated with pgACC GABA+/Cr concentrations and RSFC in women with PPD. In an exploratory aim, we examined correlations between RSFC and cortical OCC GABA+/Cr concentrations in postpartum women.

## MATERIALS AND METHODS

### Subject selection

English-speaking pregnant participants were consented and assessed for eligibility in our medical center obstetric clinics using the Edinburgh Postnatal Depression Scale (EPDS) [29]. 88 eligible and interested women between 19-40 years old consented to the prospective study (Figure 1). Two groups enrolled during pregnancy: (1) 35 healthy comparison women (HCW) and (2) 53 women at risk for PPD (AR-PPD). Details and rationale for the inclusion/exclusion criteria for both groups are in Supplementary Material. Institutional Review Boards (IRB) approved the study which was conducted between April 2013 and February 2017.

**Figure 1:**
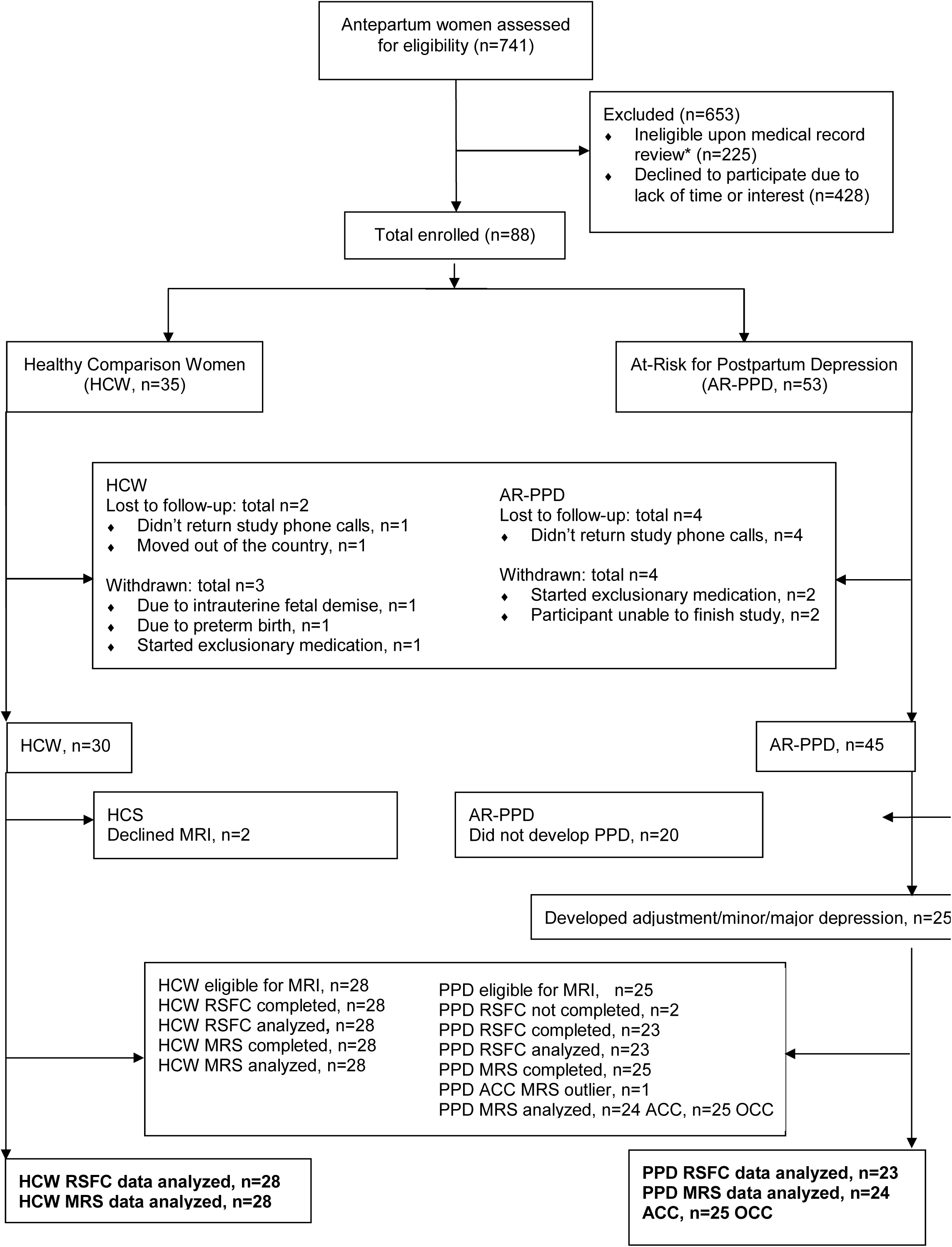
Enrollment and study completion flow diagram Footnote:

^*^ Based on medical record review, participants were excluded if they had any significant current medical illness. Available clinical laboratory results including blood cell counts, oral glucose tolerance testing, chemistry panels, thyroid function tests and viral serology conducted for routine care were reviewed and were within normal limits.

### Study procedures

Antepartum visits (visits 1 and 2) were completed in second and third trimesters. Postpartum visits were completed within days after delivery (visit 3), 2-7 weeks (visit 4) and 4-11 weeks postpartum (visit 5). HCW (n=28) and women with PPD (n=25) completed imaging (Figure 1) at either visit 4 or 5, prior to 8 weeks postpartum, based on literature suggesting an optimal postpartum onset definition[30]. The Structured Clinical Interview for DSM-IV TR Disorders (SCID-IV), Patient Edition [31], Structured Interview Guide for Hamilton Depression Rating Scale (HAM-D17) [32], Structured Interview Guide for Hamilton Anxiety Scale (HAM-A)[33], Spielberger State-Trait Anxiety Inventory (STAI)[34], Pittsburgh Sleep Quality Index (PSQI) [35], Sheehan Disability Scale (SDS) [36], urine benzodiazepine and pregnancy test were completed as detailed in Supplementary Material. Blood sample collection/handling, rationale for the timing of postpartum MRI and inclusion of women with a new onset adjustment disorder with depressed mood are delineated in Supplementary Material.

### Magnetic resonance neuroimaging procedures

### fMRI data acquisition and analysis

#### Image acquisition

Neuroimaging data was acquired with one research-dedicated 3.0T Philips Achieva whole-body MR system using phased-array receiver SENSE head coil (Philips Health-Care, Best, the Netherlands) at the Advanced MRI Center, University of Massachusetts Medical School. Image acquisition followed the same procedure as previously published[8] and is detailed in Supplementary Material.

#### fMRI data analysis

We completed both an independent components analysis (ICA) and a seed-based analysis to examine group differences in RSFC (Aims 1, 3 + 4). We conducted conventional preprocessing (motion correction, slice-timing correction and spatial smoothing (FWHM=5mm), but no temporal filtering or distortion correction) followed by ICA-AROMA to detect and regress out movement-related signal[37]. We then carried out model-free ICA using FSL MELODIC (including temporal filtering with a 150-second cut-off; 2 mm sampling) with a preset number of independent components (=30). Results of this step identified six components that contained regions in the ACC/MPFC. These components (numbered 4,6 and 8-11 out of 30), included two representing the DMN, one SN and one central executive network, and right and left frontoparietal networks; together they accounted 21.3% of explained variance and 13.42% of total variance. FSL’s dual regression and randomise procedures were then used to interrogate these six components, while the other 24 components were not further investigated. A group-differences analysis (i.e. the design matrix contained regressors for group means, age and number of postpartum days at time of MRI; randomise number of permutations=1000) was used to identify regions of significant group differences in network connectivity (TFCE *p*-corrected outputs thresholded at *p*>0.995833, representing a Bonferroni correction of 0.05/12 (6 components ^*^ 2 contrasts). This analysis identified a region in the DMPFC in one of the DMN components. We then ran two correlation analyses with dual regression, using either total HAM-D_17_ or EPDS scores as the covariate. Focusing on the single ICA component for the DMN, and thresholding at *p*>0.975 (0.05/2=0.025), both of these analyses identified essentially the same DMPFC region as found for the group-differences analysis, and no others. Because both depression-score regions were larger than the group-differences region, we chose to threshold the HAM-D_17_-correlation analysis at a slightly more stringent level (*p*>0.99), producing a DMPFC region now entirely contained within the group-differences region. This HAM-D_17_-generated DMPFC region was henceforth used as the seed region for next-step seed-based analyses, including correlation analyses with cortical GABA/Cr+ (see results). For seed-based analyses, participant-specific residual signal was extracted after regressing out white-matter and CSF signals, and was correlated with the seed region’s time-course to create each participant’s correlation map.

#### ^*1*^*H MRS data acquisition, processing and quantification*

T1-weighted anatomical MRI were collected for diagnostic and localization purposes as noted above. Edited MRS spectra were acquired using MEscher-GArwood Point-REsolved Spectroscopy Sequence (MEGA-PRESS) (TE=68 msec; TR=2000 msec; 16 averages x 20 dynamic scans)[38] (Aims 2, 3 + 4). Two MRS sequences were completed with each participant with the first voxel of interest positioned in the bilateral pgACC (3.0 cm (anterior-posterior) X 3.0 cm (left-right) X 2.0 cm (cranio-caudal)) region (Aim 2) and the second voxel of interest positioned in the midline OCC (3.0 cm (anterior-posterior) X 3.0 cm (left-right) X 3.0 cm (cranio-caudal) (Supplementary Figure 1, Validation aim). Voxel placement is described in Supplementary Materials.

Data processing and spectral fitting were done by using MATLAB (The Mathworks, MA, USA) software. MEGA-PRESS[39], a frequency selective technique, enabled us to detect the GABA+ peak of the spectra by eliminating the signal due to Cr. Prior to Fournier transformation of the MRS signal, 3 Hz exponential line broadening was applied. The difference-edited GABA+ signal at 3ppm and the Cr peak from unsuppressed PRESS signal were fit with Gaussian and Lorentzian models respectively (*r*>0.95) and the area under these model peaks was calculated to obtain GABA+ to Cr (GABA+/Cr) ratios[40]. We calculated the location and partial volume makeup of each MRS voxel and corrected for partial volume effects and relaxation as detailed in Supplementary Material. GABA spectra quality was assessed and an outlier analysis performed as detailed in Supplementary Material.

#### Plasma neuroactive steroid quantification

Antepartum and postpartum plasma concentrations of progesterone, allopregnanolone (i.e. 3α, 5α-tetrahydroprogesterone) and pregnanolone (i.e. 3α, 5β-tetrahydroprogesterone) were determined by a liquid chromatography-tandem mass spectrometry (LC-MS/MS) assay as previously published[25] with a lower limit of quantification of 0.005ng/mL, 0.008ng/mL and 0.020ng/mL respectively (Aim 4).

### Statistical Analysis

#### Demographic and clinical data

Baseline and postpartum characteristics and peripartum depression and anxiety ratings were compared between PPD vs. HCW groups using Fisher’s exact test for categorical variables and Student’s independent samples *t*-test for continuous variables (with Satterthwaite adjustment for unequal variances, when appropriate) (Aims 1-4).

#### Resting-state functional connectivity

Participant-specific seed correlation maps (for the HAM-D_17_-correlated DMPFC region) were submitted to group analyses (group-differences or correlations) using FSL Feat. Predictors included group, HAM-D_17_ or EPDS scores, and GABA+/Cr as regressors of interest, and age and number of postpartum days at time of MRI as nuisance regressors. Results of these analyses were thresholded at *z*=3.1 (*p*<.001) to identify significant voxels, and then at a cluster criterion of *p*<0.05 to qualify the resulting clusters (via Gaussian Random Field theory) (Aims 1, 3 + 4).

#### Magnetic resonance spectroscopy

For all analyses, the threshold for statistical significance (α) was 0.05. We used independent-sample t-tests to compare OCC (Validation Aim) and pgACC GABA+/Cr concentrations (Aims 2, 3, + 4) between groups and bivariate correlation to test for relationships with HAM-D_17_, EPDS and allopregnanolone. A Bonferroni correction was applied for multiple comparisons of a priori hypothesized correlations between GABA and HAM-D_17_ and EPDS as well as RSFC. These analyses were conducted in IBM SPSS Statistics for Windows, Version 24.0 (IBM Corp., Armonk, NY, USA.).

#### Peripartum plasma neuroactive steroid concentrations and analyses of relationships with postpartum mood, RSFC and GABA MRS

We analyzed the longitudinal relationship between HAM-D_17_, EPDS and plasma allopregnanolone and its isomer pregnanolone across peripartum visits (Aim 4). We also measured their precursor progesterone, since if progesterone concentrations differed between groups, it could affect metabolite (i.e. allopregnanolone and pregnanolone) concentrations.

To test for relationships between peripartum NAS concentration and RSFC, we averaged antepartum allopregnanolone or pregnanolone concentrations (visit 1 and visit 2) and separately averaged postpartum concentrations (visits 3, 4 and 5) so we could examine correlations of *antepartum* vs. *postpartum* NAS with RSFC and cortical GABA+/Cr. We correlated these average values against the functional connectivity maps for the DMPFC seed. Statistical modeling was done as previously published [25] with full details in Supplementary Material.

## RESULTS

### Demographic and clinical data

Participants were on average 28.8 (±4.9) years old, with most participants of Caucasian race (69%) and non-Hispanic ethnicity (80%) (Table 1). All peripartum psychometric scale scores significantly differed by risk group, with women diagnosed with PPD by postpartum SCID having higher scores on all scales, at all visits (Supplementary Table 1) (Aims 1-4). The HAM-D_17_ and EPDS depression scale total scores were highly correlated with each other across all participants (*r* = +.911, *p*<0.000).

**Table 1:** Participant Characteristics at Antepartum Study Entry and in the Postpartum

| Variable<sup>1</sup> | HCW<br>(n=28) | PPD<br>(n=23) | P-value<sup>2</sup> |
| --- | --- | --- | --- |
| **Antepartum Data** |  |  |  |
| **Age in years, mean (SD)** | 29.0 (5.0) | 28.6 (4.9) | 0.78 |
| **Right-handed, %** | 82.1 | 91.3 | 0.99 |
| **Race, %** |  |  |  |
| Caucasian | 60.7 | 73.9 | 0.60 |
| Black or African American | 17.9 | 8.7 |  |
| Asian | 7.1 | 0.0 |  |
| Some other race | 10.7 | 13.0 |  |
| **Ethnicity, %** |  |  |  |
| Not Hispanic or Latino | 75.0 | 82.6 | 0.74 |
| **Education level, %** |  |  |  |
| High school diploma or less | 10.7 | 13.0 | 0.48 |
| Partial or completed college | 60.7 | 73.9 |  |
| Graduate/professional degree | 28.6 | 13.0 |  |
| **Marital status, %** |  |  |  |
| Never married | 25.0 | 34.8 | 0.04 |
| Currently Married | 75.0 | 47.8 |  |
| Divorced/separated/widowed | 0.0 | 17.4 |  |
| **Employment status (currently employed), %** | 89.3 | 69.6 | 0.15 |
| **Number of children in household** |  |  |  |
| 0 | 53.6 | 43.5 | 0.43 |
| 1 | 42.9 | 39.1 |  |
| 2+ | 3.6 | 17.4 |  |
| **Household income, %** |  |  |  |
| Below \$19,000 | 17.9 | 39.1 | 0.26 |
| \$19,000-52,000 | 25.0 | 30.4 | |
| \$53,000-111,000 | 28.6 | 17.4 | |
| \$112,000 and above | 28.6 | 13.0 | |
| **Receiving financial or housing assistance from government, %** | 7.1 | 21.7 | 0.22 |
| **Planned pregnancy, %** | 67.9 | 52.2 | 0.39 |
| **Parity, %** |  |  |  |
| Nulliparous | 57.1 | 56.5 | 0.43 |
| Primiparous | 39.3 | 30.4 |  |
| Multiparous | 3.6 | 13.0 |  |
| **Thyroid Stimulating Hormone, mean (SD) in uIU/mL** | 2.1 (1.8) | 1.7 (0.6) | 0.38* |
| **Hamilton Depression Rating Scale (HAM-D<sub>17</sub>), mean (SD)** | 3.5 (2.7) | 14.1 (6.1) | <.0001* |
| **Hamilton Anxiety Rating Scale (HAM-A), mean (SD)** | 6.1 (4.2) | 18.1 (7.7) | <.0001* |
| **Edinburgh Postnatal Depression Scale, mean (SD)** | 2.2 (2.5) | 12.8 (4.0) | <.0001* |
| **Spielberger State-Trait Anxiety Inventory-State (STAI-S), mean (SD)** | 24.8 (8.1) | 44.2 (10.0) | <.0001 |
| **Sheehan Disability Scale (SDS), mean (SD)** | 1.4 (3.4) | 11.7 (6.1) | <.0001* |
| **SCID diagnosis at study entry, %** |  |  |  |
| Adjustment disorder with depressed mood | 0.0 | 4.3 | 0.45 |
| Depressive disorder, not otherwise specified | 0.0 | 56.5 | <.0001 |
| Anxiety disorder | 0.0 | 78.3 | <.0001 |
| Past depressive disorder | 0.0 | 82.6 | <.0001 |
| Past postpartum depressive disorder | 0.0 | 26.1 | 0.006 |
| **Postpartum Data** |  |  |  |
| **Gestational age at delivery (weeks), mean (SD)** | 39.4 (1.3) | 39.0 (1.2) | 0.34 |
| **Preterm delivery, %** | 3.6 | 4.5 | 0.99 |
| **Delivery method, %** |  |  |  |
| Vaginal | 85.7 | 63.6 | 0.10 |
| Cesarean section | 14.3 | 36.4 |  |
| **Labor induction, %** | 57.1 | 54.5 | 0.99 |
| **Infant weight (grams), mean (SD) at delivery** | 3258.9 (523.2) | 3496.5 (549.3) | 0.13 |
| **Infant admitted to neonatal intensive care unit, %** | 7.1 | 4.5 | 0.99 |
| **Maternal complications during labor and delivery, %** |  |  |  |
| Gestational onset hypertension | 3.4 | 4.2 | 0.99 |
| Pre-eclampsia | 6.9 | 12.5 | 0.65 |
| Postpartum hemorrhage | 0.0 | 0.0 | - |
| Premature rupture of membranes | 6.9 | 0.0 | 0.49 |
| **Breastfeeding status, %** |  |  |  |
| Full or partial | 82.1 | 81.8 | 0.99 |
| Bottle-feeding | 17.9 | 18.2 |  |
| **Thyroid Stimulating Hormone, mean (SD) in uIU/mL** | 1.5 (1.0) | 1.1 (0.5) | 0.13* |
| **Hamilton Depression Rating Scale (HAM-D<sub>17</sub>) at time of MRI, mean (SD)** | 3.2 (3.1) | 15.9 (6.5) | <.0001* |
| **Number of postpartum days at time of MRI, mean (SD)** | 38.6 (17.8) | 29.9 (12.9) | 0.05 |
| **Hamilton Anxiety Rating Scale (HAM-A) at time of MRI, mean (SD)** | 3.9 (3.6) | 18.1 (7.5) | <.0001* |
| **Edinburgh Postnatal Depression Scale at time of MRI, mean (SD)** | 1.9 (2.5) | 13.7 (3.7) | <.0001 |
| **Spielberger State-Trait Anxiety Inventory-State (STAI-S) at time of MRI, mean (SD)** | 25.4 (7.8) | 47.9 (10.3) | <.0001 |
| **Sheehan Disability Scale (SDS) at time of MRI, mean (SD)** | 0.9 (2.6) | 12.0 (6.4) | <.0001 |
| **SCID diagnosis at time of MRI, %** |  |  |  |
| Major depression with peripartum onset | 0.0 | 43.5 | <.0001 |
| Minor depression with peripartum onset | 0.0 | 39.1 | 0.0003 |
| Adjustment disorder with depressed mood | 0.0 | 17.4 | 0.04 |
| Anxiety disorder | 0.0 | 73.9 | <.0001 |

<sup>†</sup> Unless otherwise indicated, data are expressed as number (percentage) of subjects. Percentages have been rounded and may not total 100.
<sup>2</sup>P-values from Fisher's Exact Test for categorical variables and Student's independent samples t-test for continuous variables (with Satterthwaite adjustment for unequal variances, when appropriate, designated by \*).
SD: Standard Deviation; PPD: Postpartum depression; HCW: Healthy comparison women

### Resting-state functional connectivity analyses

Within the DMN network identified with ICA, dual-regression analyses identified a significant group difference in the DMPFC (peak voxel: MNI coordinates (2, 58, 32) *p*=0.002), (Figure 2). This region demonstrated greater connectivity with the rest of the DMN in PPD as compared to HCW (Aim 1 +4). We also conducted two correlation analyses (across all participants, again using dual-regression) which indicated that both HAM-D_17_ (Figure 2) and EPDS total scores were positively correlated with RSFC values in an overlapping DMPFC region (peak HAM-D_17_ voxel: MNI coordinates (0, 60, 34), *p*=0.008; peak EPDS voxel: MNI coordinates (0, 60, 34), p=0.004) (Aim 1 + 4).

**Figure 2:**
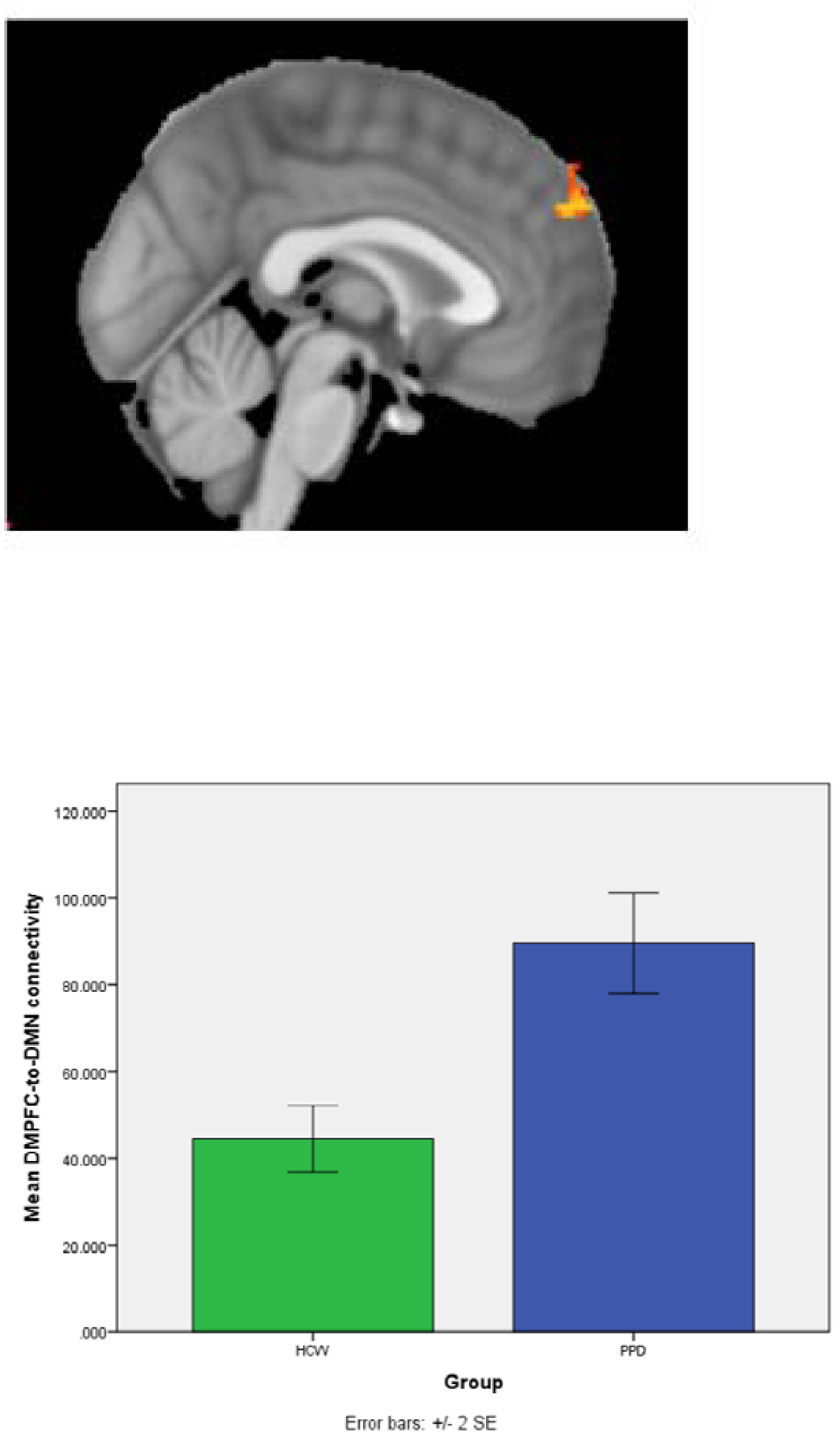
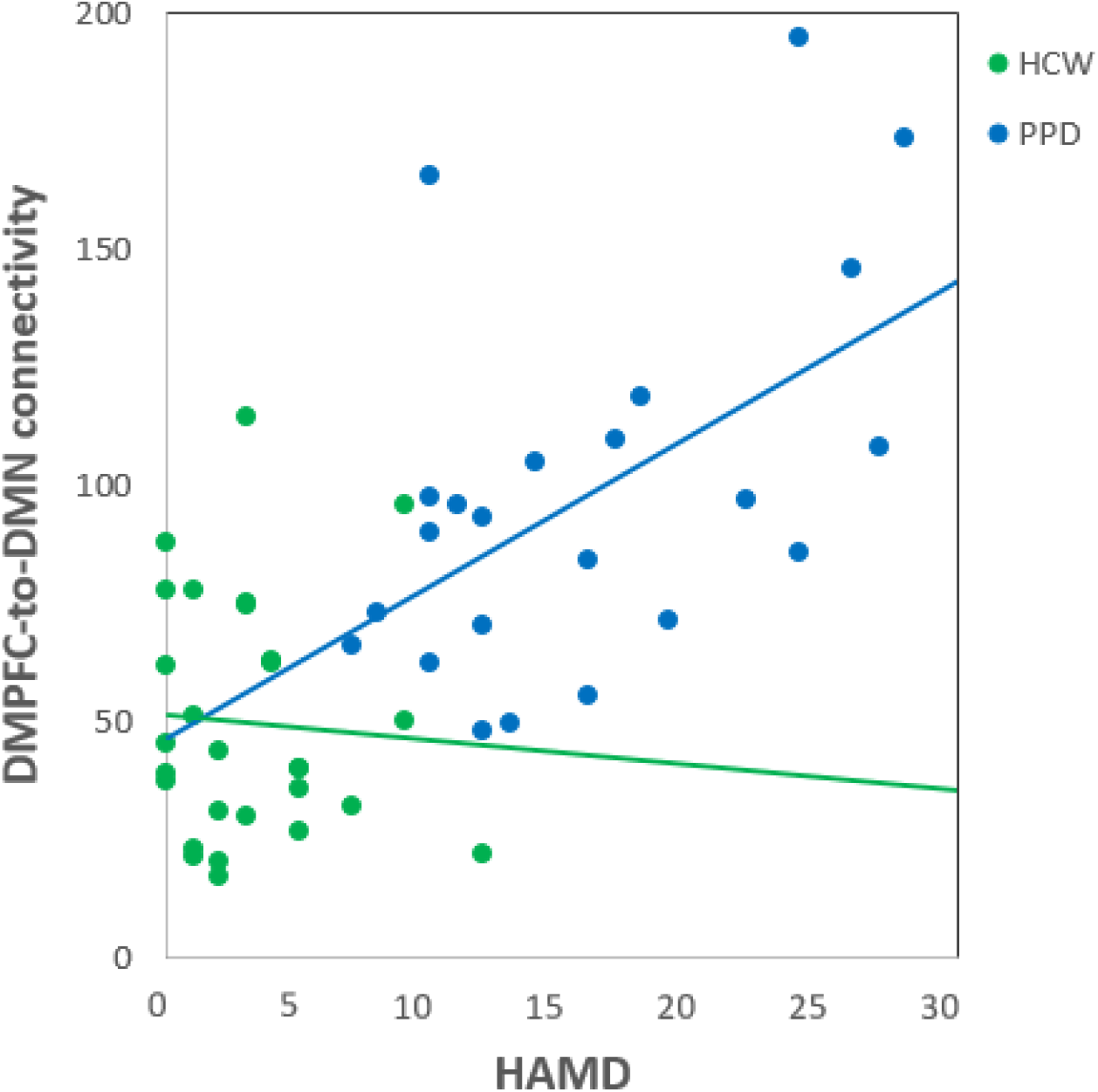
ICA analysis identified a default-mode network (DMN) component where connectivity of dorsomedial prefrontal cortex (DMPFC) to the rest of the network was related to peripartum depression (PPD). The DMPFC cluster with greater resting-state functional connectivity in PPD as compared to healthy comparison women (peak voxel: MNI coordinates (2, 58, 32) is (across all postpartum participants) positively correlated with HAM-D_17_ scores (HAM-D_17_ peak voxel: MNI coordinates (0, 60, 34).

Mean functional connectivity with the DMPFC seed across all participants is shown in Supplementary Figure 2. Supplementary Table 2 lists regions where significant group differences in connectivity with the DMPFC seed were found: HCW had stronger connectivity in the precuneus, posterior cingulate and postcentral gyrus and supramarginal gyrus/angular gyrus regions (Aims 1 + 4). None of these regions with group differences in DMPFC connectivity correlated with either HAM-D_17_ or EPDS total scores (*p*>0.05).

### ^1^H-MRS GABA+/Cr concentrations in the pgACC and OCC

Fifty-three participants (PPD, n=25, HCW, n=28) completed the MRS scan (Figure 1). Cr concentrations were measured in both regions of interest and there were no between-group differences (data not shown). The mean pgACC GABA+/Cr concentration in HCW and PPD was 0.10369(*±*0.03907) and 0.106213 (±0.07069) units, respectively. The mean OCC GABA+/Cr concentration in HCW and PPD was 0.108914 (*±*0.01293) and 0.106122 (*±*0.01211) units, respectively. Neither the pgACC nor OCC GABA+/Cr concentrations differed between groups (both *p*> 0.4). GABA+/Cr concentrations were not correlated with either HAM-D_17_ or EPDS, either across all participants, or separately within groups (all *p*> 0.3) (Aims 2 + Validation).

### Resting-state functional connectivity and ^1^H-MRS GABA+/Cr correlation analyses

GABA+/Cr measures were correlated with whole-brain resting-state DMPFC connectivity, controlling for age and total postpartum days, and using a voxel-wise threshold of *z*=3.0 and cluster threshold of *p*=.05 (Aim 3). For all regions so identified (Supplementary Table 3), mean z-scores were extracted and correlated against each individual’s pgACC and OCC GABA+/Cr. Across all participants, pgACC GABA+/Cr correlated positively with DMPFC RSFC in a region spanning the right anterior/posterior insula as well as the right temporal pole (r=+0.661, p=0.000; Supplementary Table 3; Figure 3). Within this region, extracted values for the PPD group alone showed a correlation of *r*=+0.546, *p*=0.013 with postpartum pgACC GABA+/Cr, while the HCW group showed a correlation of *r*=+0.700, *p*=0.000. Using the same whole-brain approach, OCC GABA+/Cr correlated positively with regions spanning both amygdalae (right amygdala: *r*=+0.522, *p*=0.000; left amygdala: *r*=+0.651, *p*=0.000; Supplementary Table 3; Figure 3). Post-hoc tests revealed that the PPD group alone showed correlations with right amygdala (*r*=+0.547, *p*=0.010) and left amygdala (*r*=+0.569, *p*=0.007) while the HCW group alone showed correlations with right amygdala (*r*=+0.445, *p*=0.023) and left amygdala (*r*=0.724, *p*=0.000. OCC GABA+/Cr measures also correlated positively with parietal areas (all participants: *r*=+0.532, *p*=0.000; PPD alone: *r*=+0.524, *p*=0.015; HCW alone: *r*=+0.560, *p*=0.003), and negatively with a region spanning supplementary motor cortex and dorsal ACC (all participants: *r*=-0.530, *p*=0.000; PPD alone: *r*=-0.397, *p*=0.075; HCW alone *r*=-0.605, *p*=0.001; Supplementary Table 3; Figure 3). All post-hoc tests were uncorrected.

**Figure 3:**
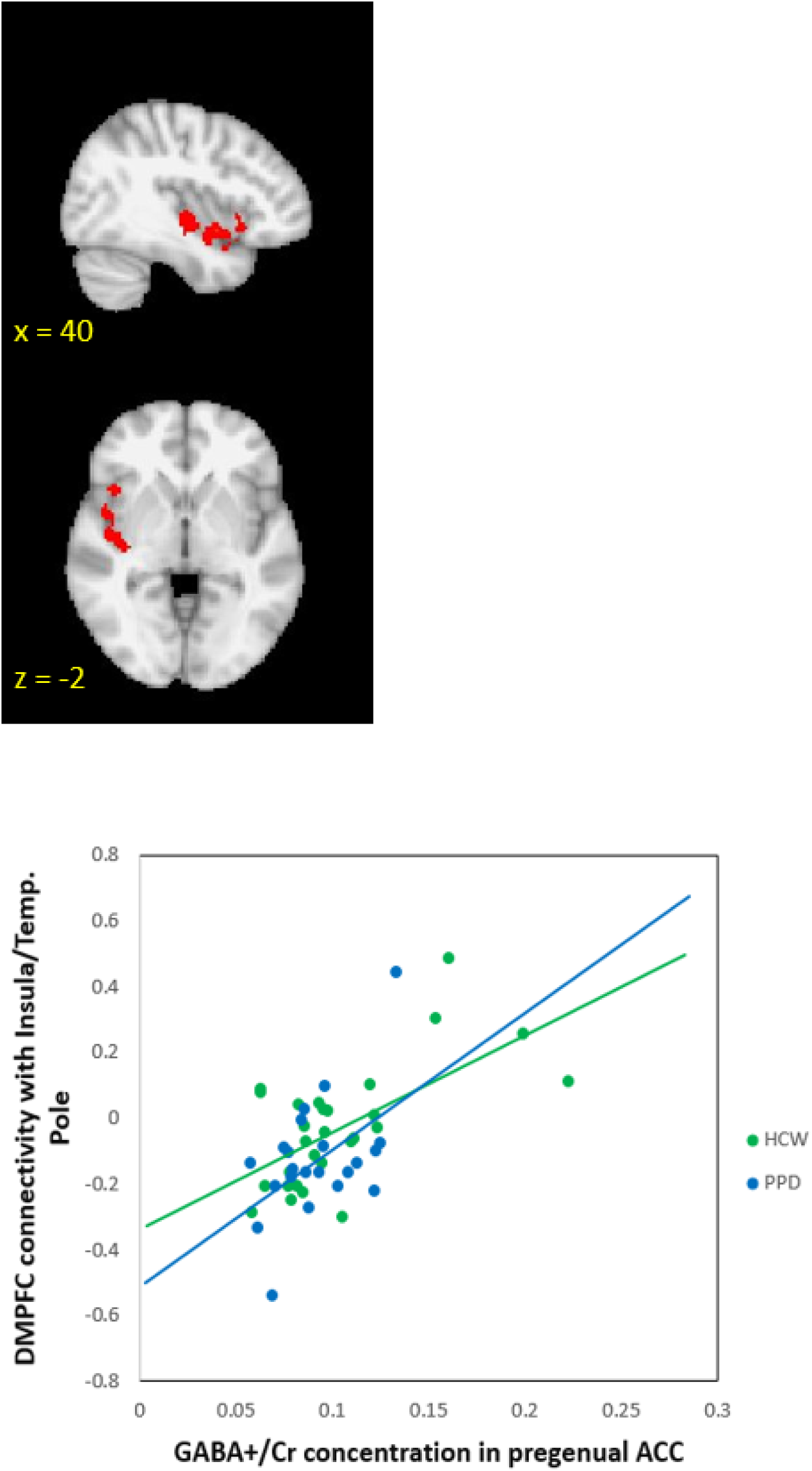
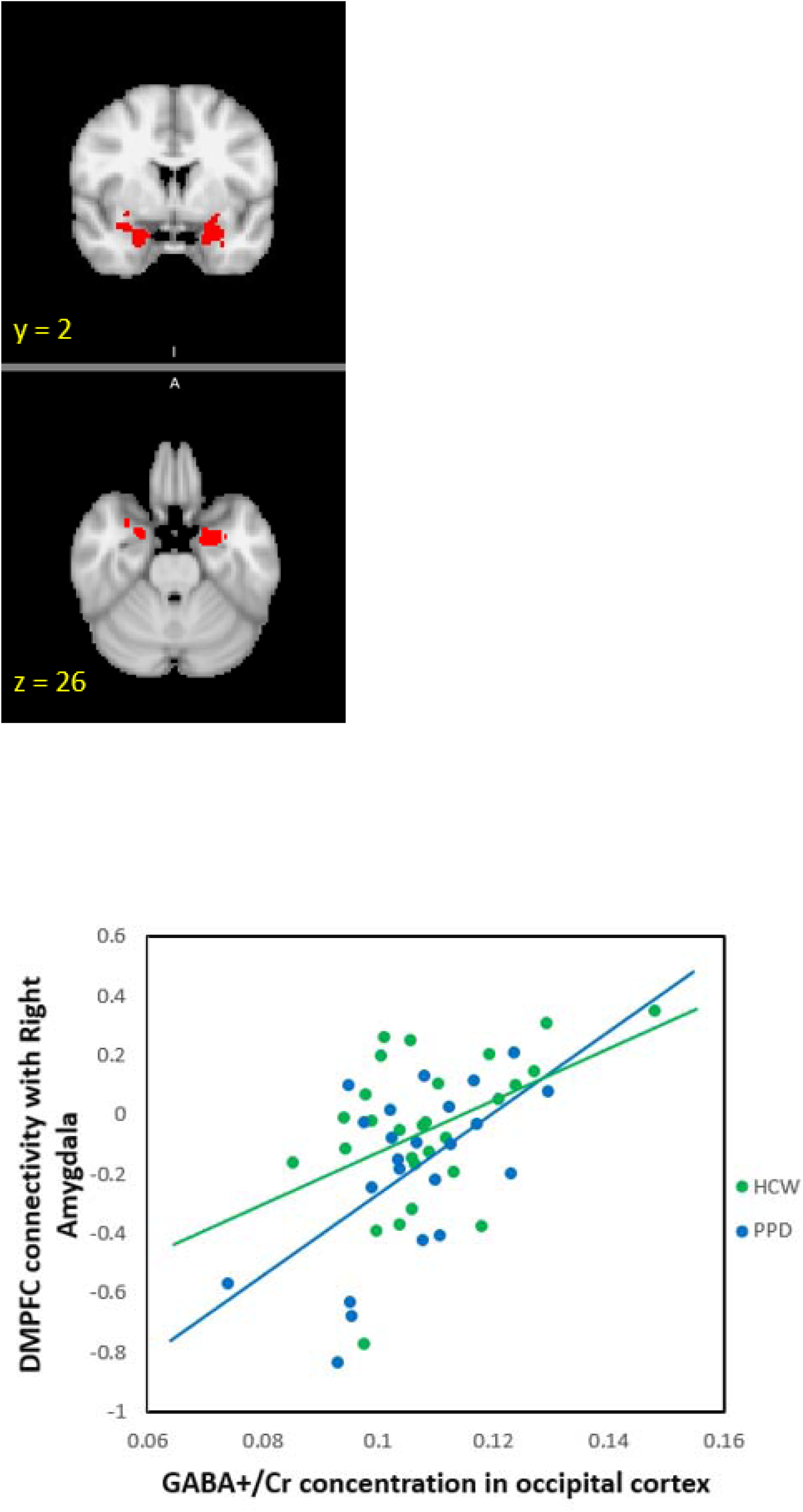
Brain regions where dorsomedial prefrontal cortex (DMPFC) resting-state connectivity is correlated with ^1^H-MRS GABA+/Cr concentrations across postpartum participants.

[UPPER] Figure depicts a region spanning the insula and temporal pole where DMPFC resting-state functional connectivity is significantly correlated with pregenual anterior cingulate cortex GABA+/Cr concentration. Scatterplot shows relationship with insula/temporal pole.

[LOWER] Figure depicts areas in the right and left amygdalae where DMPFC resting-state functional connectivity is significantly correlated with occipital cortex GABA+/Cr concentration. Scatterplot shows relationship in right amygdala.

Abbreviations: ^1^H-MRS = proton magnetic resonance spectroscopy; GABA = gamma-aminobutyric acid; Cr= creatine

### Peripartum plasma neuroactive steroid concentrations and relationship to postpartum mood, resting-state functional connectivity and GABA MRS

Antepartum, peripartum and postpartum progesterone and pregnanolone concentrations did not significantly differ between groups (data not shown). *Peripartum* allopregnanolone was 0.97 +/- 0.42 ng/mL higher in women with PPD (*p*=0.03) (Aim 4).

Main effects models demonstrated that HAM-D_17_ and EPDS were marginally associated with greater *peripartum* allopregnanolone (β=0.05 +/- 0.02, *p*=0.03 for HAM-D_17_; β=0.05 +/- 0.03, *p*=0.05 for EPDS). Allopregnanolone concentrations were correlated against DMPFC connectivity using the same whole-brain thresholds (voxel z>3.0; cluster *p*<=.05, controlling for age and postpartum days), as described for GABA. Across participants, averaged *postpartum* allopregnanolone concentrations were significantly correlated with intra-DMPFC connectivity (*r*=+0.548, *p*=0.000; Figure 4). For the PPD group alone, the correlation was significant for the PPD group (*r*=+0.794, *p*=0.000), but not for the HCW group (*p*>0.29). By contrast, *antepartum* allopregnanolone concentrations were not significantly correlated with any region in the DMPFC maps for either group or across all participants. Likewise, either for separate groups, or across all participants, neither *antepartum* nor *postpartum* pregnanolone concentrations were significantly correlated with RSFC in the DMPFC maps. There were no significant correlations between the averaged antepartum or postpartum allopregnanolone or pregnanolone concentrations and OCC or pgACC 1H MRS GABA+/Cr concentrations (Aim 4).

**Figure 4:**
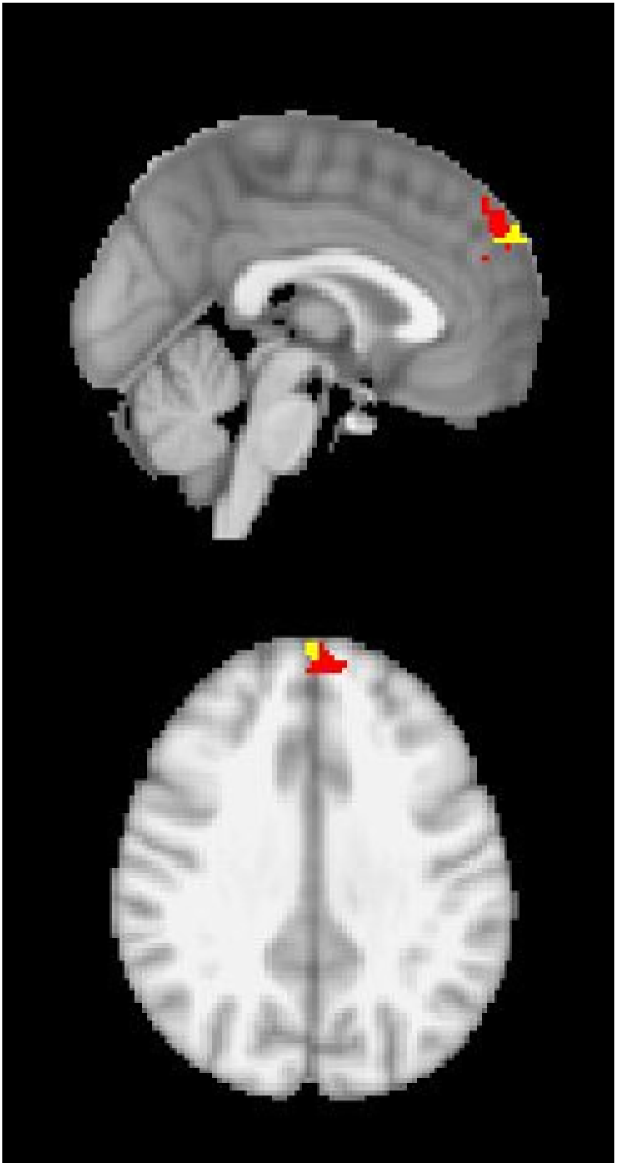
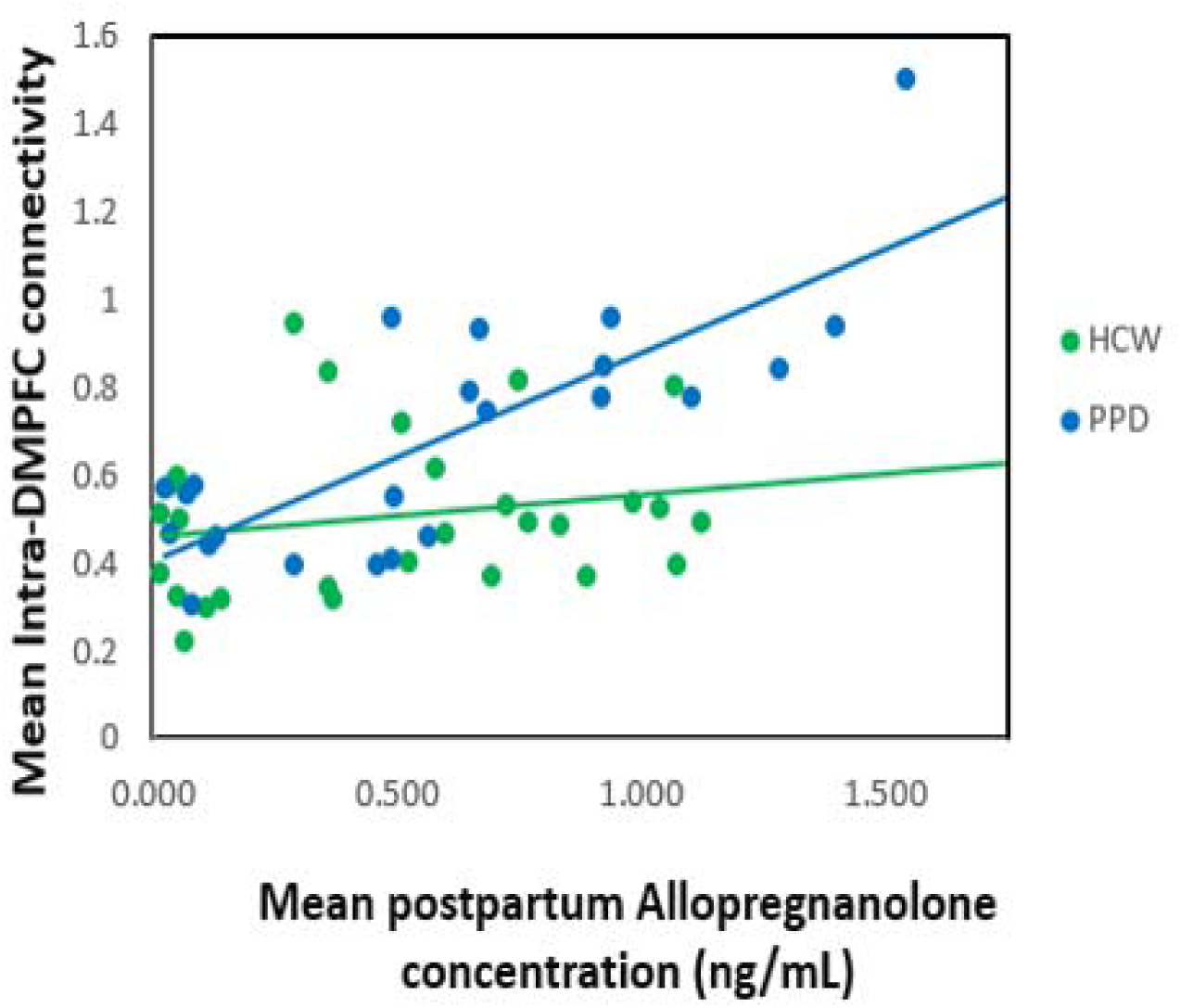
Region in the dorsomedial prefrontal cortex (red) where sconnectivity values (z-scores) are significantly related to average plasma postpartum allopregnanolone concentrations across all postpartum participants. (The small yellow region shows the area of significant correlation between HAMD scores and the default-network component identified in the ICA analysis.) The allopregnanolone-DMPFC correlation is significant even when data for the outlier subject is excluded.

Abbreviations: HAMD: Hamilton Depression Rating Scale; ICA: Independent Component Analysis

## DISCUSSION

This study measured plasma NAS, RSFC using fMRI and cortical GABA+/Cr concentrations using ^1^H-MRS in women who developed PPD during gestation up to 8 weeks after delivery as compared to healthy peripartum women. Our preliminary findings suggest that the relationship between allopregnanolone and GABA and RSFC within the DMN and SN is important in the pathophysiology of PPD. An area of the DMPFC has greater connectivity with the rest of the DMN in women with peripartum adjustment disorder with depressed mood or minor/major depressive disorder (Aim 1). Across all postpartum women, depression severity was significantly positively correlated with RSFC within this DMPFC (Aim 4). Greater RSFC between the DMN component and this DMPFC mirrors reports of RSFC abnormalities between the MPFC and DMN regions in MDD[41]. We identified reduced RSFC between the DMPFC and the precuneus, posterior cingulate cortex and supramarginal gyrus/angular gyrus regions in PPD. The precuneus is structurally connected to the posterior cingulate cortex, the ACC, thalamus, and striatum[42] and has a role in self-related mental representations during rest and first-person perspective taking[43]. The posterior cingulate cortex, seated at the nexus of multiple intrinsic connectivity networks, including the DMN, has a central role in internally directed cognition[11]. The supramarginal gyrus is important for emotion perception[44]. Taken together, altered connectivity between the DMPFC and these regions could manifest clinically as a disturbance of self-appraisal and emotional perception in PPD.

We anticipated early postpartum cortical GABA concentrations to be low in comparison to those reported during the follicular or luteal phase [17,18] and similar to postpartum concentrations previously reported[18]. In rodents, region-specific changes in GABAergic inhibition occur in the peripartum period[45]. Elevated antepartum NAS are associated with inhibition of glutamic acid decarboxylase (GAD) mRNA expression, reducing brain GABA concentrations[46] while the postpartum is characterized by reduced allopregnanolone and enhanced neuronal GABA synthesis, raising brain GABA concentrations in regions important to maternal care[47]. Contrary to our hypothesis, we did not identify a between-group difference in GABA+/Cr (Aim 2). We designed the study to include a heterogeneous at-risk population. However, since the underlying mechanisms for each clinical phenotype may differ, inclusion as a group might have obscured the ability to identify biological relationships. Inclusion of adjustment disorder with depressed mood may have reduced our ability to identify potential between-group differences, as we are not aware of evidence for reduced GABA in adjustment disorder. Alternatively PPD may differ from MDD which has been associated with reduced cortical GABA concentrations[16].

pgACC and OCC GABA+/Cr concentrations were associated with intrinsic connectivity of the DMPFC in postpartum women regardless of mood (Aim 3 + Validation). Across participants, pgACC GABA+/Cr correlated positively with RSFC in a region spanning the right anterior/posterior insula as well as the right temporal pole. Additionally, across participants OCC GABA+/Cr concentrations correlated positively with regions spanning both amygdalae. Some of the regions where we identified DMPFC RSFC correlations with cortical GABA+/Cr concentrations are considered within the SN. Our preliminary results demonstrate that RSFC in some regions of the SN have a positive relationship with GABA+/Cr in postpartum women. It is unknown if this association exists in non-postpartum women. Ranges of OCC GABA+/Cr (both PPD and HCW) and ACC GABA+/Cr (HCW only) are narrow, but the range of ACC GABA+/Cr in PPD is wider. We believe the wider range of ACC GABA+/Cr concentrations reflects the range measured in postpartum depressed and anxious women, rather than due to MRS technique or due to known factors which affect cortical GABA levels.

Contrary to our hypothesis, peripartum plasma allopregnanolone concentration was *higher* in women who developed PPD as compared to HCW and marginally associated with depression scores (Aim 4). There was no significant between-group difference in allopregnanolone’s isomer, pregnanolone, nor was it correlated to RSFC or GABA MRS measures. We based our hypothesis on our previous report of increased 5-β reduction of progesterone to pregnanolone in peripartum women at-risk for PPD[25] compared to HCW, a study that did not include women with PPD.

Postpartum plasma allopregnanolone was positively correlated with postpartum intra-DMPFC connectivity. Contrary to our hypothesis, we did not identify a correlation between plasma allopregnanolone and cortical GABA+/Cr concentrations in postpartum women (Aim 4). This finding is in agreement with the previous MRS study[18], but is unexpected given the role of NAS in GABA neurochemistry as earlier reviewed. Acute stress can increase brain allopregnanolone levels[48] and this may or may not be reflected in the peripheral levels measured. We believe our data lends support to the hypothesis that peripartum allopregnanolone is associated with intrinsic functional connectivity and may contribute to the development of PPD. The differences we report in peripartum allopregnanolone concentrations could represent differential metabolism within the progesterone-based biosynthetic pathway, either peripherally and/or in the brain[21]. Altered peripartum allopregnanolone concentrations could affect cortical GABA concentrations and affect downstream GABAergic inhibitory tone within brain regions we did not investigate.

There are several strengths of the study design. Inclusion/exclusion criteria were stringent to maximize rigor. We utilized both research-clinician administered instruments as well as self-report measures to make our findings relevant to both research and clinical settings. The longitudinal design allowed for repeated mood and NAS assessment and for imaging to be completed before women with moderate PPD initiated pharmacotherapy. An additional strength is the use of an ultra-sensitive LC-MS/MS assay[25].

Due to our study design, we cannot distinguish between women who had PPD onset in pregnancy vs. early postpartum as we conducted the SCID at antepartum study visit 1 and at the postpartum imaging session (visit 4 or 5). It is possible that there are connectivity and neurochemistry differences between women with different times of onset. Our results apply to postpartum women meeting diagnostic criteria for major and minor depressive disorder and adjustment disorder with depressed mood: importantly, more than 70% of these women had a comorbid anxiety disorder. Although we pre-screened a diverse population, women who participated were mainly white, non-Hispanic women, which could limit the generalizability of the results. In order to obtain valid results, our study excluded women with additional important risk factors for PPD including psychotropic medication use, recent substance abuse and active medical disorders. The study includes a moderate sample size. It would have been helpful to include a healthy non-pregnant follicular phase population for comparison. Finally, we report “GABA+” to acknowledge that our measured concentration includes the presence of macromolecules[49] which may be considered a limitation.

In conclusion, we report preliminary evidence of relationships between allopregnanolone and intrinsic functional connectivity in the DMN in PPD, and cortical GABA+/Cr and RSFC in postpartum women. It is possible that peripartum allopregnanolone, through positive allosteric modulatory effects on GABA, contributes to the differences we see in DMN connectivity in PPD. These data are significant as we can now further investigate how postpartum neurochemistry and circuitry may differ from the non-postpartum state allowing us to investigate postpartum-specific therapeutic targets.

## ACKNOWLEDGEMENTS

We sincerely thank Dr. Peter J. Schmidt for his mentorship throughout the career development award and for his feedback on the manuscript. We additionally thank Dr. Todd Lencz for his thoughtful comments during data analysis and preparation of the manuscript and gratefully thank the peripartum women who participated in this research. The functional connectivity results were previously presented in a poster at the American College of Neuropsychopharmacology (ACNP) 56^th^ Annual Meeting, Palm Springs, CA, December, 2017.

## SUPPLEMENTARY MATERIALS

Supplementary Information accompanies this paper at https://doi.org/- doi to be determined.

## LEGENDS FOR TABLES AND FIGURES

**SUPPLEMENTAL FIGURE 1:** Proton magnetic resonance spectroscopy (^1^H-MRS) voxel localization and ^1^H spectra.

Footnote: Upper part illustrates voxel localization for the occipital cortex (left) and pregenual anterior cingulate cortex (right) superimposed onto a slice from the anatomical scan of a healthy comparison woman. The lower part is a sample representation of an edited ^1^H spectrum for a healthy comparison woman. The blue line indicates the unedited ^1^H spectrum, the green line indicates the edited ^1^H spectrum and the red line indicates an attenuated version of the green line.

Abbreviations: MNI: Montreal Neurological Institute; HAM-D_17_: Hamilton Depression Rating Scale; EPDS: Edinburgh Postnatal Depression Scale

**SUPPLEMENTAL FIGURE 2:** Mean correlation, across all participants, of the dorsomedial prefrontal cortex (DMPFC) seed. Significant functional coupling is displayed by voxels ranging from red to yellow for positive correlations.

**SUPPLEMENTAL TABLE 1:** Peripartum Psychometric Scale Total Scores (mean ±SD)

**SUPPLEMENTAL TABLE 2:** Regions where resting-state functional connectivity with the dorsomedial prefrontal cortex (DMPFC) seed region differs significantly by group. Footnote: All coordinates in the Montreal Neurological Institute (MNI) atlas space. Peaks whose coordinates have greater than 50% probability of lying in white matter are not listed Abbreviations: Post. = posterior; G. = gyrus; Sup. = superior; L= lobule

**SUPPLEMENTAL TABLE 3:** Brain regions where dorsomedial prefrontal cortex (DMPFC) functional connectivity is correlated with pregenual anterior cingulate cortex (pgACC) or occipital cortex (OCC) ^1^H-MRS GABA+/Cr concentrations

